# Endothelial VCAM1 links oncogenic KRAS signaling to myeloid inflammation and hemorrhagic instability in brain arteriovenous malformations

**DOI:** 10.64898/2026.11.15.766869

**Authors:** Jung-Eun Park, Hyejin Park, Ohnmar Myint, Sung Wook Kang, Xian Chen, Zhongming Zhao, Jongmin Choi, Sung Yun Jung, Shuning Huang, Jakob Körbelin, Huimahn Alex Choi, Jaroslaw Aronowski, Peng Roc Chen, Eunhee Kim, Eun S. Park

**Affiliations:** Vivian L. Smith Department of Neurosurgery, McGovern Medical School, The University of Texas Health Science Center at Houston, Houston, TX 77030, USA; Center for Neuroimmunology and Glial Biology, The Brown Foundation Institute of Molecular Medicine, The University of Texas Health Science Center at Houston, Houston, TX 77030, USA; Michael E. DeBakey Department of Surgery, David J. Sugarbaker Division of Thoracic Surgery, Baylor College of Medicine, Houston, TX 77030, USA; Center for Precision Health, School of Biomedical Informatics, The University of Texas Health Science Center at Houston, Houston, TX, 77030, USA; Lester and Sue Smith Breast Center, Baylor College of Medicine, Houston, TX 77030, USA; The Verna and Marrs McLean Department of Biochemistry and Molecular Pharmacology, Baylor College of Medicine, Houston, TX, 77030, USA; Department of Diagnostic and Interventional Imaging, McGovern Medical School, The University of Texas Health Science Center at Houston, Houston, TX 77030, USA; Department of Oncology, Hematology and Bone Marrow Transplantation, University Medical Center Hamburg-Eppendorf, 20246 Hamburg, Germany; Department of Neurology, McGovern Medical School, The University of Texas Health Science Center at Houston, Houston, TX 77030, USA

**Keywords:** Brain arteriovenous malformation, Intracerebral hemorrhage, KRAS mutation, Endothelial cells, VCAM1, Microglia, Macrophages, Inflammation

## Abstract

**BACKGROUND:** Hemorrhagic conversion is the most devastating complication of brain arteriovenous malformations (bAVMs). However, endothelial mechanisms promoting macrophages (Mϕ) recruitment, microglia (MG) activation, and bAVM lesion instability remain unclear. We investigated vascular cell adhesion molecule-1 (VCAM1) as a mediator linking oncogenic endothelial signaling to myeloid cell responses and hemorrhagic progression in bAVMs.

**METHODS:** We performed integrated transcriptomic analysis of human bAVM and epilepsy-control brain tissues, with validation in local bAVM specimens. Transcriptomic and proteomic analyses of bAVM lesions and intact brain tissues from mutant KRAS-induced bAVM mice were followed by histological validation. Using AAV-BR1-KRAS^G12V^-injected *Cx3cr1-gfp;Ccr2-rfp* mice, we performed intravital imaging to assess Mϕ/MG responses to VCAM1 neutralization. Hemorrhage and myeloid cell responses were also evaluated following *endothelial-specific VCAM1 deletion*. Plasma soluble VCAM1 (sVCAM1) was measured in patients with unruptured and ruptured bAVMs and unruptured aneurysm controls. Effects of sVCAM1 on MG activation were examined *in vitro*.

**RESULTS:** Integrated human endothelial bulk and single-cell RNA-sequencing and tissue validation identified VCAM1 upregulation in bAVMs, including unruptured lesions. Plasma analyses showed increased sVCAM1 in unruptured and ruptured bAVM patients compared with unruptured aneurysm controls. In mutant KRAS-induced bAVM mice, transcriptomic and proteomic analyses demonstrated increased VCAM1 expression, with endothelial localization confirmed histologically. *In vitro,* mutant KRAS increased sVCAM1 production, while recombinant sVCAM1 stimulated inflammatory mediator expression in MG. Both MG and Mϕ expressed VLA-4, a major VCAM1 receptor. Intravital imaging showed that VCAM1 neutralization reduced Mϕ accumulation and MG activation, while early VCAM1 neutralization reduced hemorrhagic burden. Consistently, *endothelial-specific VCAM1 deletion* reduced Mϕ accumulation, MG activation, and hemorrhage volume in bAVM mice and improved behavioral performance.

**CONCLUSIONS:** These findings identify endothelial VCAM1 as an upstream mediator linking mutant KRAS signaling to myeloid cell responses and hemorrhagic progression in bAVMs. This study establishes endothelial VCAM1 as a potential therapeutic target for attenuating hemorrhagic progression.

Graphical Abstract:
This study identifies endothelial VCAM1 as an upstream mediator of the inflammatory vascular response in brain arteriovenous malformation (bAVM), contributing to macrophage recruitment, microglial activation, and hemorrhagic progression. Integrated analyses of human transcriptomic datasets, bAVM tissues, and patient plasma samples revealed increased endothelial VCAM1 expression and elevated circulating soluble VCAM1 (sVCAM1). In a mutant KRAS-driven mouse model of bAVM, transcriptomic and proteomic profiling, together with pharmacological VCAM1 neutralization and endothelial-specific VCAM1 deletion, demonstrated that endothelial VCAM1 contributes to myeloid cell accumulation, microglial responses, and hemorrhagic progression. Collectively, these findings support endothelial VCAM1 as a potential therapeutic target to attenuate myeloid cell responses and vascular inflammation, thereby attenuating bAVM instability and hemorrhagic progression. Created in BioRender.

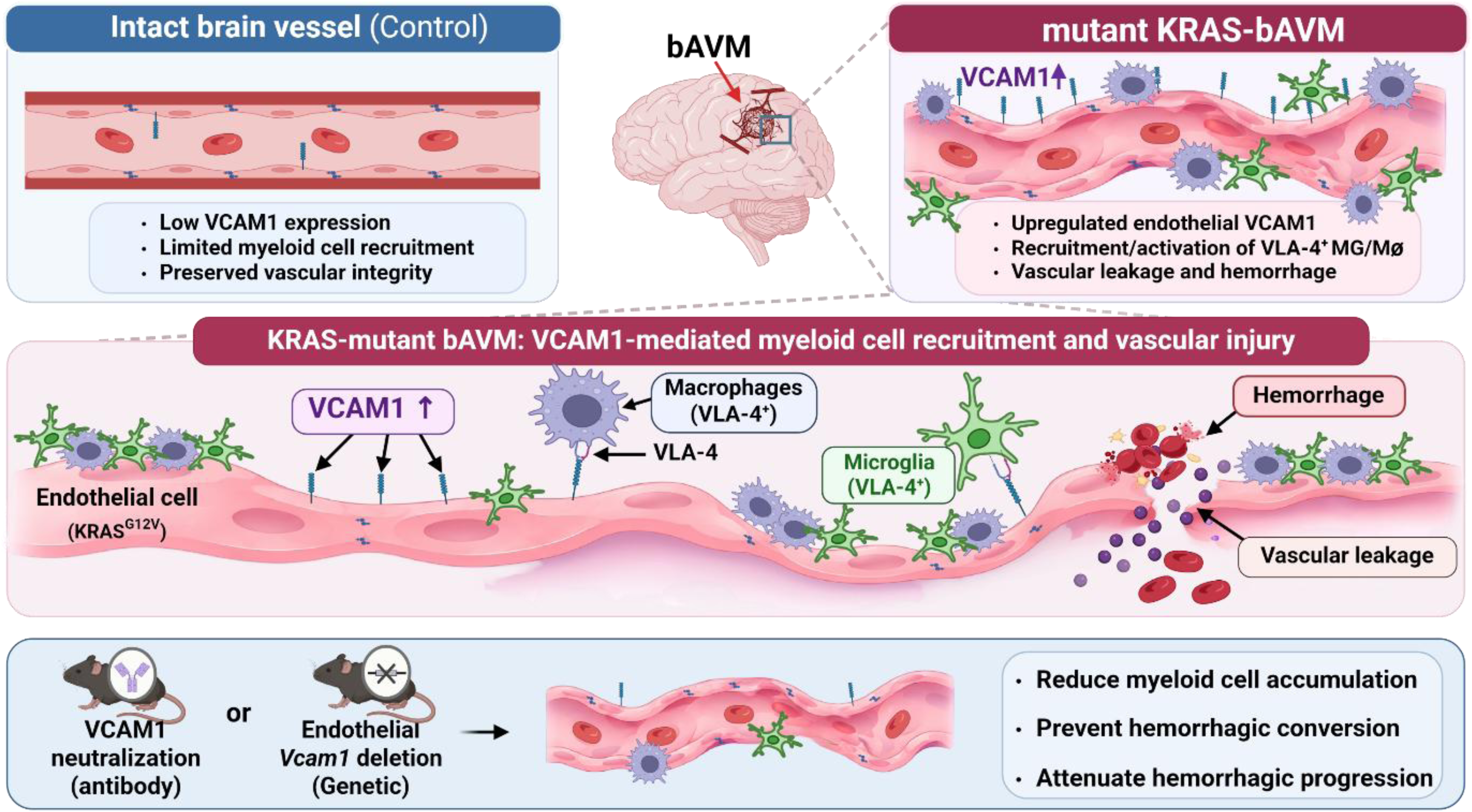

**Novelty and Significance:** *What is Known?:*

- Brain arteriovenous malformations (bAVMs) are a major cause of hemorrhagic stroke and neurological disability or death. Somatic activating KRAS mutations in endothelial cells drive abnormal angiogenic signaling and are detected in more than 72% of sporadic bAVMs.
- Microglia/macrophages (MG/Mϕ) accumulate in unruptured human and mouse bAVMs. Our previous studies demonstrated that MG/Mϕ-mediated vascular inflammation contributes to blood-brain barrier (BBB) disruption, bAVM instability, and hemorrhagic conversion in mouse bAVMs.
- VCAM1-VLA-4 signaling is an established mechanism of myeloid cell adhesion and recruitment, including macrophage recruitment, however, whether endothelial VCAM1 contributes to inflammatory bAVM instability and hemorrhagic conversion in bAVMs remains unknown.

*What new information does this article contribute?:*

- Endothelial VCAM1 is identified as an upstream mediator linking oncogenic KRAS signaling to inflammatory MG/Mϕ responses in bAVMs. Integrated human and mouse transcriptomic and proteomic analyses, together with histological validation, identify endothelial VCAM1 as a conserved and markedly upregulated molecular feature of bAVMs.
- Endothelial-derived soluble VCAM1 (sVCAM1) provides a potential mechanism for endothelial-microglial communication. Plasma sVCAM1 levels were elevated in patients with unruptured bAVMs compared with unruptured aneurysm controls, with a trend toward further elevation in ruptured bAVMs. Mutant KRAS increased endothelial sVCAM1 production, and sVCAM1 stimulated microglial activation *in vitro*, identifying a potential mechanism by which mutant endothelial cells promote brain-resident immune cell activation.
- Endothelial VCAM1 contributes functionally to myeloid cell responses and hemorrhagic instability in bAVMs. Pharmacological VCAM1 neutralization reduced Mϕ infiltration, MG activation, and early bleeding in mouse bAVMs, whereas endothelial-specific VCAM1 deletion reduced Mϕ infiltration, MG activation, and hemorrhage volume. These complementary loss-of-function approaches establish endothelial VCAM1 as a functional mediator of inflammatory bAVM instability.

**Summary of Novelty and Significance:** Hemorrhagic conversion is the most devastating complication of bAVMs, yet the endothelial mechanisms that initiate inflammatory responses and promote bAVM lesion instability remain poorly understood. Because early Mϕ infiltration and MG activation contribute to BBB disruption and hemorrhagic conversion, identifying the upstream endothelial signals that drive these responses may reveal new approaches to stabilize bAVMs. Through cross-validation of independent human transcriptomics datasets and orthogonal analyses of human bAVM tissues, we identified endothelial VCAM1 as a consistently elevated molecular feature of bAVMs. Integrated transcriptomic and proteomic analyses, *in vivo* imaging, pharmacological neutralization, and endothelial-specific genetic loss-of-function studies further demonstrate that VCAM1 links oncogenic KRAS signaling to Mϕ recruitment, MG activation, and hemorrhagic instability. In addition, our findings identify endothelial-derived sVCAM1 as a potential paracrine signal capable of activating MG.

Together, these findings establish endothelial VCAM1 as a functional mediator of the inflammatory axis connecting oncogenic vascular signaling to myeloid cell responses and hemorrhagic instability in bAVMs. This study identifies VCAM1 as a potential therapeutic target for limiting hemorrhagic progression and provide a mechanistic framework for understanding how oncogenic endothelial signaling contributes to inflammatory vascular dysfunction beyond bAVMs.

## INTRODUCTION

Brain arteriovenous malformations (bAVMs) can cause intracerebral hemorrhage (ICH), which accounts for 40-70% of deaths among affected patients.^1–3^ Moreover, bAVM-associated ICH is the most common cause of hemorrhagic stroke in children.^4^ Understanding the mechanisms underlying hemorrhagic conversion of bAVMs is therefore essential for developing therapies to reduce hemorrhagic risk. Surgical resection remains an important treatment for bAVMs; however, invasive intervention can cause substantial neurological morbidity, particularly when bAVM lesions are located in eloquent brain regions. In addition, bAVM recurrence remains a clinically important concern after apparently complete surgical obliteration, particularly in children and young patients presenting with hemorrhage, with reported 5-year recurrence rates of approximately 18-21%.^5,6^ These limitations highlight the need for therapeutic strategies that prevent bAVM instability and hemorrhagic conversion. Because bAVMs consist of direct arteriovenous connections lacking an intervening microcapillary bed and contain multiple vascular and inflammatory cell populations,^7^ identifying cellular pathways that promote bAVM instability remains an important unmet need.

Recent studies show that somatic activating mutations in KRAS, including p.Gly12Val and p.Gly12Asp, occur in endothelial cells (EC) of more than 72% of sporadic bAVM patients and are enriched in hemorrhagic bAVM lesions.^8–10^ Mutant endothelial KRAS has been established as a causal driver of bAVM formation using brain EC-targeted adeno-associated virus (AAV) vectors based on the BR1 capsid and genetic mouse models.^11,12^ In our previous studies, we established that endothelial KRAS activation is sufficient to promote the recruitment of peripheral immune cells, including monocyte-derived macrophages (Mϕ), and activation of brain-resident microglia (MG) in a mutant KRAS-driven bAVM mouse model.^11^ Increased myeloid cell accumulation has similarly been observed in hemorrhagic lesions of human bAVMs and mouse models of hereditary hemorrhagic telangiectasia (HHT).^13,14^ We further demonstrated that Mϕ and MG promote vascular inflammation, impair endothelial junction integrity, and disrupt the blood-brain barrier (BBB), thereby contributing to hemorrhagic conversion in bAVMs. However, how mutant KRAS signaling in ECs initiates Mϕ recruitment and MG activation in the bAVM area remains poorly understood.

Recruitment of circulating immune cells into the brain is critically regulated by endothelial adhesion molecules.^15^ Vascular cell adhesion molecule-1 (VCAM1) mediates adhesion or transmigration of peripheral immune cell such as monocyte-derived Mϕ, leukocyte, or neutrophils, particularly through interactions with its major integrin receptor, very late antigen-4 (VLA-4; α4β1, encoded by ITGA4 and ITGB1).^16–18^ VLA-4 binding to endothelial VCAM1 promotes firm immune cell adhesion and subsequent transendothelial migration, providing a potential mechanism through which endothelial activation regulates inflammatory cell recruitment. Notably, increased endothelial VCAM1 expression and myeloid cell expression of VLA-4 have been associated with inflammatory cell recruitment and vascular injury in hemorrhagic stroke.^19,20^ In bAVMs, increased endothelial VCAM1 expression has also been reported in human and experimental mouse bAVM lesions.^21,22^ However, whether endothelial VCAM1 actively promotes myeloid cell recruitment and MG activation, thereby contributing to hemorrhagic progression in bAVMs, remains largely unknown. Given our previous findings that early Mϕ recruitment and MG activation in unruptured bAVMs before hemorrhagic conversion in mutant KRAS-induced bAVMs, determining whether endothelial VCAM1 initiates these inflammatory responses may reveal an upstream mechanism underlying bAVM instability.

In this study, integrated analyses of human transcriptomic datasets, local bAVM tissues, and plasma samples, together with transcriptomic and proteomic profiling of mutant KRAS-induced mouse bAVMs, identified VCAM1 as a prominent endothelial adhesion molecule associated with bAVM pathology. We found that endothelial VCAM1 was consistently increased in unruptured human bAVMs and mutant KRAS-induced mouse bAVMs, accompanied by Mϕ infiltration and MG activation. Using intravital imaging, endothelial-specific VCAM1 deletion, and *in vitro* studies, we demonstrated that oncogenic KRAS signaling induces endothelial VCAM1, which promotes the recruitment of VLA-4-expressing Mϕ and contributes to MG activation, resulting in hemorrhagic instability in bAVMs. Together, these findings identify endothelial VCAM1 as a potential mechanistic link between oncogenic KRAS signaling, myeloid cell recruitment, and inflammatory vascular instability, providing new insights into the mechanisms underlying hemorrhagic progression in bAVMs.

## METHODS

The Supplemental Methods section can be found in the Supplemental Material.

## RESULTS

### Human bAVMs exhibit endothelial VCAM1 upregulation in unruptured bAVM lesions

We previously observed accumulation of MG/Mϕ in human bAVM tissues,^23^ however the endothelial mechanisms promoting Mϕ recruitment and MG activation remain poorly understood. To identify candidate endothelial pathways involved in these processes, we re-analyzed bulk transcriptomic data from ECs isolated from human bAVM tissues and compared them with ECs isolated from the unaffected cortex of patients with epilepsy.^24^ Among the upregulated genes, we identified two major endothelial adhesion molecules, VCAM1 and intercellular adhesion molecule-1 (ICAM1). Both VCAM1 and ICAM1 were significantly increased in bAVM ECs, with VCAM1 exhibiting a greater magnitude of upregulation and higher discriminatory capacity than ICAM1 (VCAM1: logFC = 1.46, AUC = 0.756; ICAM1: logFC = 0.99, AUC = 0.722) (**Figure 1A** through **1C**).

**Figure 1.**
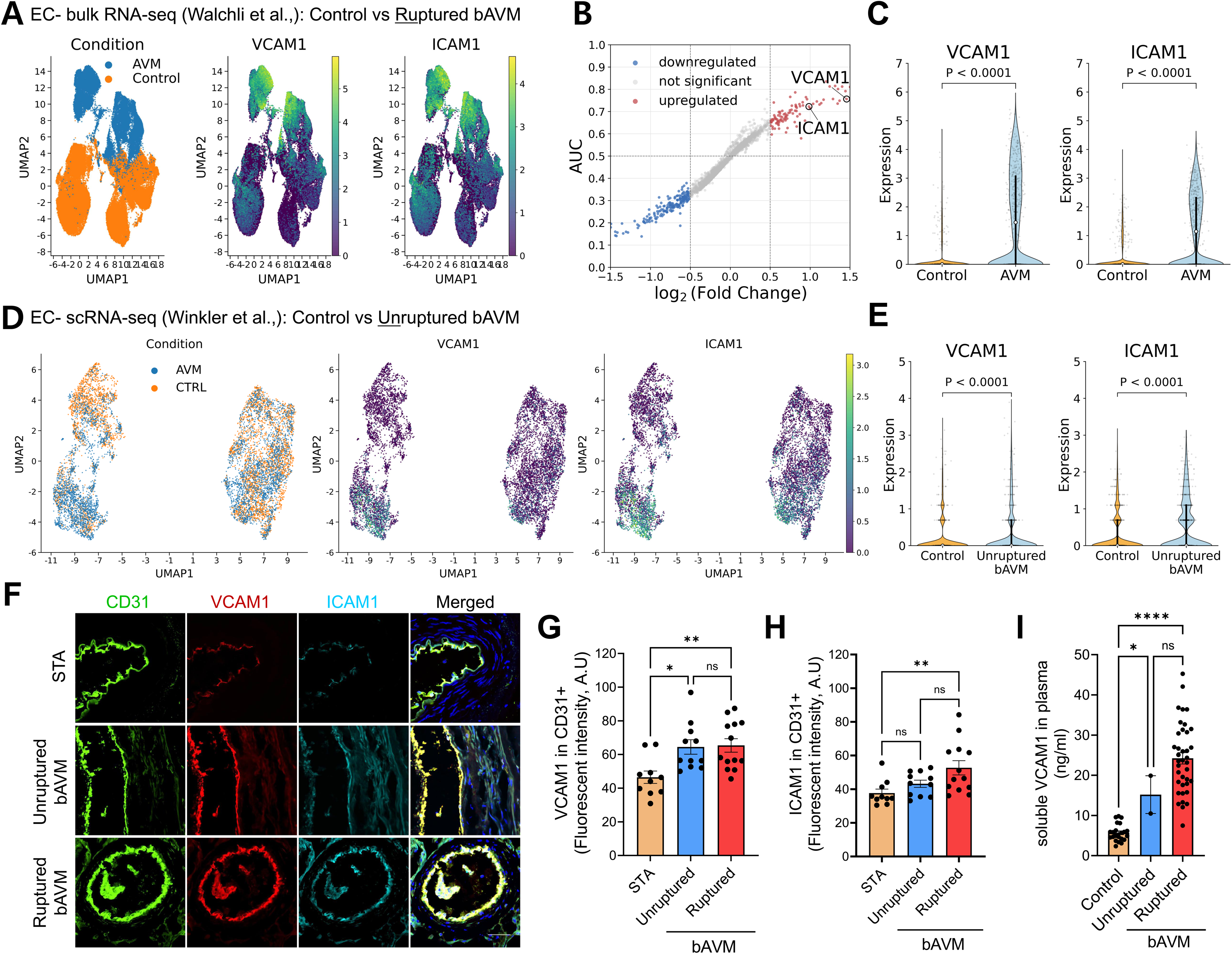
Human bAVMs exhibit increased endothelial VCAM1 expression. **A** through **C,** Re-analysis of human ruptured bAVM and control tissue bulk RNA-seq data from Walchli et al., 2024.^24^ (bAVM, n=5 donors; epilepsy control cortex, n=3 donors). **A.** UMAP visualization of ECs from control (orange) and bAVM (blue) tissues, with VCAM1 and ICAM1 expression overlaid. Each dot represents an individual EC. **B,** AUC-based differential expression analysis showing log2 fold change (log2FC) versus discriminatory effect size [(AUC-0.5) x 2). Genes were classified as upregulated (red), downregulated (blue), or not significant (gray), based on adjusted *p*<0.05 and log2FC thresholds of ≥ 0.5. VCAM1 and ICAM1 are indicated by open circles. **C,** Violin plots showing VCAM1 and ICAM1 expression in control and bAVM ECs. Dots represent a randomly selected subset of up to 500 cells per group. White circles indicate medians, and black bars indicate interquartile ranges (IQRs). Statistics significance was assessed using a two-sided Mann-Whitney U test. **D** and **E,** Re-analysis of scRNA-seq data from unruptured human bAVM tissues (n=13 donors) and control cortical vessels from epilepsy patients (n=5 donors), based on Winkler et al., 2022.^25^ **D.** UMAP visualization of ECs from control (orange) and unruptured bAVM (blue) tissues, with VCAM1 and ICAM1 expression overlaid. Each dot represents an individual cell. **E.** Violin plots showing VCAM1 and ICAM1 expression. Statistical significance was determined using a two-sided Mann-Whitney U test. **F** through **H,** Representative immunofluorescence images (**F**) and quantification of endothelial VCAM1 (**G**) and ICAM1 (**H**) fluorescence intensity in human unruptured and ruptured bAVM tissues and non-bAVM associated Superficial Temporal Artery (STA) controls. Scale bar = 50 μm. Fluorescence intensity is expressed in arbitrary units (A.U). ANOVA. *, *p*<0.05; **, *p*<0.01; ns, not significant. Each dot indicates a randomly selected region of interest (ROI; 10-13 ROIs per group) obtained from Human specimens (n=3 or 4 per group). **I,** Bar graphs quantifying sVCAM1 levels in plasma collected from ruptured, unruptured bAVM patients, compared to the plasma from unruptured aneurysm patients (n=22). Each dot indicates individual subject: ruptured (n=38), unruptured (n=2), unruptured aneurysm (n=22). ANOVA, \**p*<0.05; \*\*\*\**p*<0.0001

To determine whether endothelial adhesion molecule expression was elevated in bAVMs in the absence of overt hemorrhage, we next re-analyzed EC clusters from a published single-cell RNA-sequencing (scRNA-seq) dataset of unruptured bAVM tissues and control cortical vessels from patients with epilepsy.^25^ Both VCAM1 and ICAM1 expression were increased across EC clusters in unruptured bAVMs compared with control vessels (**Figure 1D** and **1E**), demonstrating that endothelial adhesion molecule upregulation is present in unruptured bAVM lesions.

We next validated VCAM1 and ICAM1 expression in human bAVM tissues encompassing unruptured and ruptured lesions, using the superficial temporal artery (STA) as a vascular control. Both endothelial VCAM1 and ICAM1 expression were significantly increased in ruptured bAVM tissues compared with STA controls. Notably, endothelial VCAM1 expression was also significantly elevated in unruptured bAVMs compared with STA controls (increase of 17.96 A.U), with no significant difference between unruptured and ruptured lesions. In contrast, ICAM1 expression showed a progressive increase in mean expression from STA controls to unruptured (increase of 5.54 A.U) and to ruptured bAVMs (increase of 15.0 A.U), although statistical significance was reached only for the ruptured-versus-STA comparison (**Figure 1F** through **1H**). Thus, although both adhesion molecules were elevated in unruptured bAVMs at the transcriptomic level, tissue validation demonstrated a more consistent association of VCAM1 with unruptured lesions. Given the established role of VCAM1 in myeloid cell adhesion and recruitment,^26^ these findings suggest that endothelial VCAM1 may contribute to myeloid cell recruitment in bAVMs in the absence of overt hemorrhage. Accordingly, we prioritized VCAM1 for further investigation as a potential endothelial mediator of myeloid cell recruitment and neuroinflammatory responses in bAVMs.

Meanwhile, because endothelial VCAM1 can undergo proteolytic shedding to generate soluble VCAM1 (sVCAM1),^27,28^ we next examined circulating sVCAM1 levels. Plasma sVCAM1 levels were elevated in patients with unruptured bAVMs (n=2) compared with patients with unruptured aneurysms (n=22) and were also elevated in patients with ruptured bAVMs (n=40) compared with aneurysm controls (**Figure 1I**). Although sVCAM1 levels tended to be higher in ruptured than in unruptured bAVMs, this difference was not statistically significant. Because of the limited availability of plasma samples from patients with unruptured bAVMs, this comparison should be interpreted as exploratory. Nevertheless, these findings provide preliminary evidence of increased circulating sVCAM1 in patients with bAVMs, including those without overt hemorrhage, consistent with increased VCAM1-related endothelial activity.

Together, these complementary transcriptomic, scRNA-seq, tissue, and plasma analyses identify VCAM1 as a consistently elevated endothelial adhesion molecule in human bAVMs, including unruptured bAVM lesions. The elevation of VCAM1 in unruptured bAVMs, which was also observed in ruptured lesions, suggest that VCAM1 may contribute to the inflammatory microenvironment associated with Mϕ recruitment and MG activation, potentially promoting vascular instability and hemorrhagic rupture. These findings provided the rationale for investigating endothelial VCAM1 as an upstream mediator of these inflammatory responses and hemorrhagic instability in bAVMs.

### Mutant KRAS induces endothelial VCAM1 expression *in vivo* and *in vitro*

Somatic activating KRAS mutations are detected in a substantial proportion of sporadic bAVMs.^8^ We therefore investigated whether endothelial mutant KRAS signaling is associated with VCAM1 upregulation using a mouse model of mutant KRAS-induced bAVMs.^11^ To characterize the cellular and molecular alterations associated with bAVM lesions and corresponding intact brain regions at 14 weeks after AAV-BR1-KRAS^G12V^ injection, together with proteomics profiling at 8 weeks after injection. Tissue regions were identified by T2 weighted-MRI and included established ruptured bAVM lesions and corresponding intact regions (**Figure 2A**). The scRNA-seq dataset was annotated to identify major vascular and inflammatory cell populations, including ECs. Within the EC population, VCAM1 expression was increased in bAVM compared with intact regions (**Figure 2B**).

**Figure 2.**
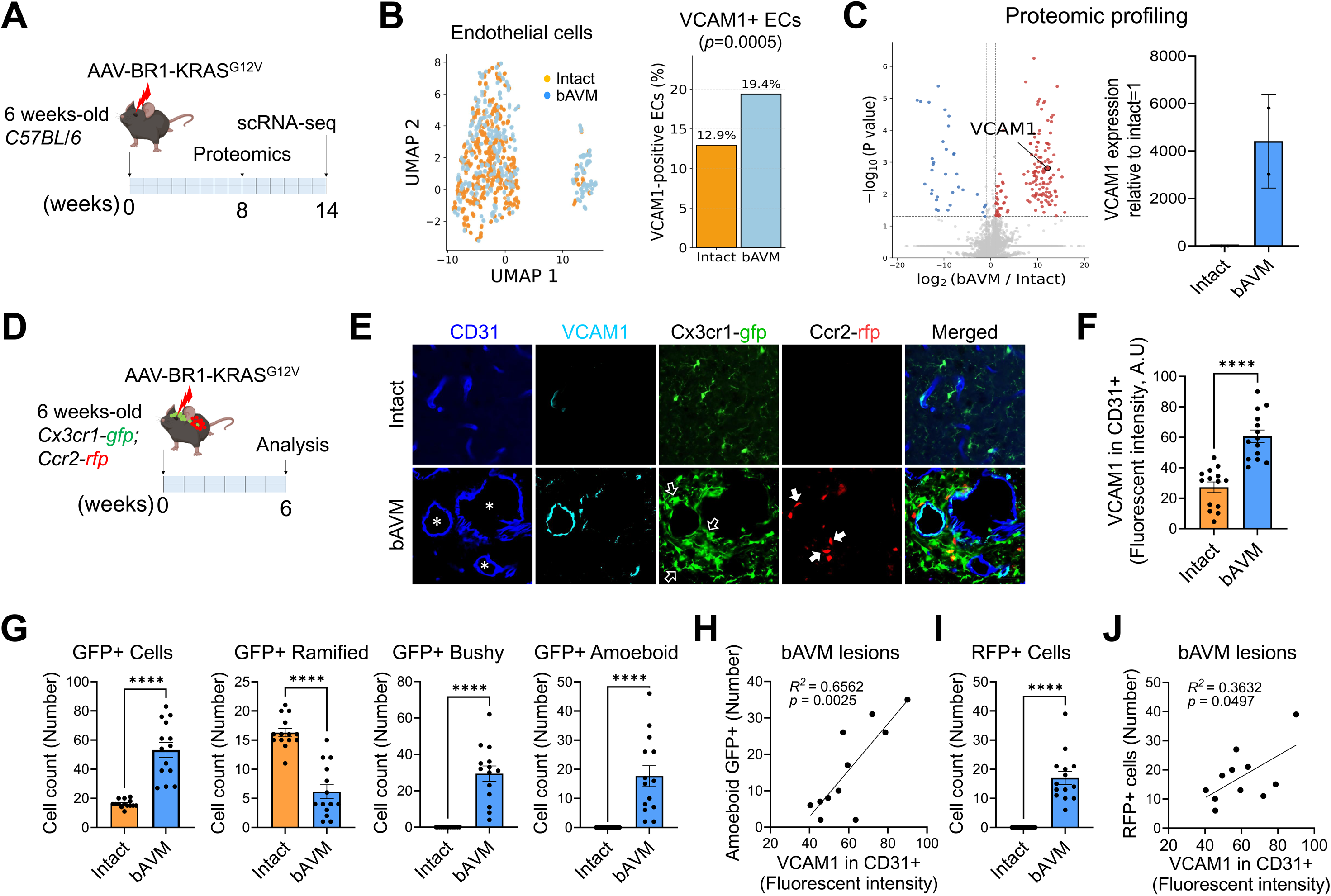
Mutant KRAS-induced bAVMs exhibit increased endothelial VCAM1 expression and myeloid cell accumulation. **A,** Schematic showing the experimental time points for scRNA-seq analysis (14 weeks) and proteomics profiling (8 weeks) after AAV-BR1-KRAS^G12V^ injection. **B,** scRNA-seq analysis of EC clusters from intact brain regions and bAVM lesions. UMAP visualization shows EC transcriptomes from Intact (orange) and bAVM (blue). Bar graph quantifies the proportion of VCAM1-expressing ECs in each group. Fisher’s exact test. **C,** Proteomic profiling of intact brain regions and bAVM lesions collected from AAV-BR1-KRAS^G12V^-injected mice at 8 weeks after injection. The volcano plot shows differentially abundant proteins, with increased VCAM1 abundance in bAVM lesions relative to intact brain tissue. The bar graph shows the relative abundance of VCAM1 in the two groups. Fisher’s exact test. n=2 mice per group. **D,** Schematic showing the experimental timeline. *Cx3cr1-gfp;Ccr2-rfp* mice received AAV-BR1-KRAS^G12V^ or -AAV-BR1-eGFP (control) and were analyzed 6 weeks after injection. **E,** Representative immunofluorescence images showing increased VCAM1 expression (cyan) in CD31-positive (+) vessels (blue), together with accumulation of CX3cr1-gfp+ MG (green) or Ccr2-rfp+ Mϕ (red) in bAVM lesions. Open arrows indicate Cx3cr1-gfp+ MG and arrows indicate Ccr2-rfp+ Mϕ. Asterisk indicate enlarged vascular lumens in bAVM lesions. Scale bar = 50 μm. **F**, Bar graphs quantify VCAM1 fluorescence intensity in CD31+ vessels from intact and bAVM lesions. Unpaired t-test. ****, *p*<0.0001. Each dot indicates a randomly selected ROI (n=14) obtained from mice (n=5) per group. **G** through **J,** Quantification of Cx3cr1-gfp+ MG for total, ramified, bushy, and amoeboid based on morphology assessment (**G**) and Ccr2-rfp+ Mϕ (**I**) in bAVM lesions compared to intact. Unpaired t-test. ****, *p*<0.0001. Each dot indicates a randomly selected ROI (n=15) obtained from mice (n=5) per group. **H, J,** Scatter plots showing the relationship between VCAM1 expression in CD31+ vessels and the abundance of GFP+ amoeboid MG (**H**) or RFP+ Mϕ (**J**) in bAVM lesions. Simple linear regression was used to assess the relationships. Each dot indicates a randomly selected ROI (n=11) obtained from mice (n=5) per group.

To determine whether increased VCAM1 abundance was detectable at an earlier stage of bAVM development, we next performed proteomics profiling at 8 weeks after AAV-BR1-KRAS^G12V^ injection. T2 weighted-MRI-guided tissue sampling was used to isolate ruptured bAVM regions, with corresponding intact regions serving as controls. Proteomic analysis revealed a marked increase in VCAM1 abundance in bAVM tissue, with a log2fold change of 12.11 relative to intact tissue (**Figure 2C**). These findings provide complementary proteomic evidence of increased VCAM1 abundance at an earlier stage of bAVM development than that examined by scRNA-seq.

We next validated VCAM1 protein expression histologically at 6 weeks after AAV-BR1-KRAS^G12V^ injection. Immunofluorescence staining demonstrated robust endothelial VCAM1 expression in bAVM vessels, whereas only low VCAM1 signal was detected in intact vessels (**Figure 2D** through **2F**). Consistent with these *in vivo* findings, mutant KRAS overexpression increased VCAM1 protein expression in cultured mouse ECs (**Figure S1A**).

We next examined the spatial relationship between endothelial VCAM1 expression and myeloid cell accumulation in bAVM lesions. Using *Cx3cr1-gfp;Ccr2-rfp* mice, in which GFP preferentially labels Cx3cr1-MG and RFP labels infiltrating monocyte-derived Ccr2-Mϕ, we observed increased VCAM1 expression accompanied by accumulation of infiltrating Mϕ and morphological activation of MG in bAVM regions **(Figure 2E**, **2G** through **2J)**. These observations are consistent with our previous findings that Mϕ/MG accumulation accompanies vascular inflammation in mutant KRAS-induced bAVMs and suggest a potential relationship between endothelial VCAM1 upregulation and local myeloid cell responses.

In addition to its membrane-bound form, VCAM1 can undergo proteolytic shedding to generate sVCAM1,^29–31^ which has been implicated in inflammatory signaling.^29^ We therefore examined whether mutant KRAS-induced endothelial VCAM1 upregulation was accompanied by increased sVCAM1 release from ECs. Conditioned medium (CM) from mutant KRAS-overexpressing ECs (KRAS-EC) contained increased sVCAM1 compared with CM from GFP-transfected control ECs (**Figure S1B**). To investigate whether sVCAM1 could directly influence microglial inflammatory responses, we treated cultured BV2-MG with recombinant sVCAM1. sVCAM1 treatment increased the expression of inflammatory and angiogenic mediators, including IL-6, IL-1β, MMP9, and VEGF-A (**Figure S1C**). Notably, these responses were similar to the inflammatory responses induced by KRAS-EC-CM in our previous study.^32^ These findings suggest that endothelial-derived sVCAM1 may contribute to microglial inflammatory activation in bAVMs.

Together, these findings demonstrate increased endothelial VCAM1 expression in mutant KRAS-induced bAVMs and support a potential role for endothelial-derived sVCAM1 in promoting microglial inflammatory responses. The spatial association of VCAM1 upregulation with Mϕ accumulation and MG activation further supports a link between endothelial VCAM1 and myeloid cell responses in bAVM lesions.

### VLA-4 is expressed by Mϕ/MG in bAVM lesions, and VCAM1 neutralization reduces myeloid cell accumulation

The above human and mouse studies provided a mechanistic rationale for investigating whether endothelial VCAM1 regulates myeloid cell recruitment, MG activation, and hemorrhagic conversion in unstable bAVMs. Because VCAM1 primarily binds very late antigen-4 (VLA-4; α4β1), a heterodimeric integrin composed of ITGA4 (α4 subunit) and ITGB1 (β1 subunit),^33^ we next examined VLA-4 expression in Mϕ and MG within bAVM lesions. Analysis of the scRNA-seq dataset from mutant KRAS-induced bAVMs revealed that both Mϕ and MG expressed VLA-4 components, with significantly increased ITGA4 expression in bAVM compared with intact regions (*p*<0.0001) (**Figure 3A** and **3B; Figure S2**). We further validated increased VLA-4 expression in Mϕ/MG in bAVM lesions compared with intact vessels at 6 weeks after AAV-BR1-KRAS^G12V^ injection (**Figure 3C** and **3D**). These findings demonstrate that VLA-4-expressing Mϕ/MG are present in bAVM lesions and provide a rationale for investigating a potential role of VCAM1-VLA-4 interactions in myeloid cells-endothelial communication.

**Figure 3.**
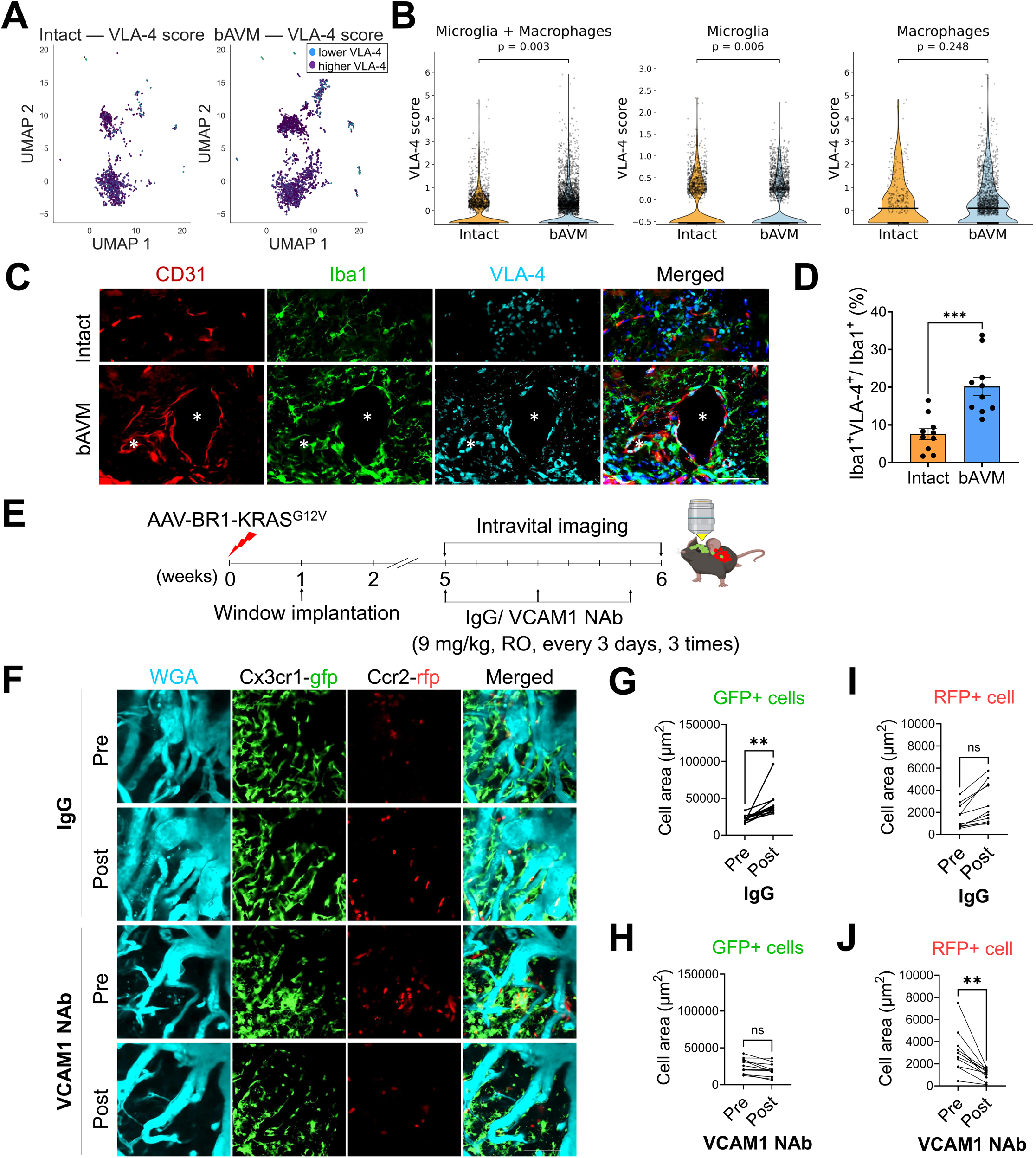
VLA-4 is expressed by MG and Mϕ in bAVM lesions, and VCAM1 neutralization reduces myeloid cell accumulation. **A.** UMAP visualization of MG and Mϕ analysis of scRNA-seq dataset from intact and bAVM lesions, classified according to lower (blue) or higher (purple) VLA-4 expression scores. **B.** Violin plots showing VLA-4 expression scores in combined MG/Mϕ populations and in MG and Mϕ populations analyzed separately. The VLA-4 score was calculated from combined ITGA4 and ITGB1 expression. Statistical significance was determined using a two-sided Mann-Whitney U test. **C,** Representative immunofluorescence images showing VLA-4 expression (cyan) in Iba1+ MG/Mϕ (green) in intact and bAVM lesions (CD31+, red). Asterisks indicates the enlarged vascular lumens in bAVM lesions. Scale bar = 50 μm. **D**, Quantification of VLA-4+ fluorescence area in Iba1+ MG/Mϕ in intact and bAVM lesions. Unpaired t-test. ***, *p*<0.001. Each dot indicates a randomly selected ROI (n=10) obtained from mice (n=3) per group. **E,** Schematic showing the cranial-window implantation and intravital imaging protocol. A cranial window was implanted 1 week after AAV-BR1-KRAS^G12V^ injection in *Cx3cr1-gfp;Ccr2-rfp* mice. Intravital imaging was performed at 5 and 6 weeks after AAV injection. **F,** Representative intravital images showing Cx3cr1-gfp+ MG and Ccr2-rfp+ Mϕ surrounding WGA-labeled dysplastic vessels in bAVM lesions. bAVM mice received control IgG or VCAM1 Neutralizing Antibody (NAb) three times at 3-day intervals, starting 5 weeks after AAV-BR1-KRAS^G12V^ injection. Images were acquired before treatment (week 5) and after treatment (week 6). Scale bars = 100 μm. **G** through **J,** Quantification of fluorescent pixel area occupied by Cx3cr1-gfp+ cells (**G**, **H**) and Ccr2-rfp+ cells (**I**, **J**) surrounding dysplastic vessels before (Week 5) and after (week 6) treatment with IgG or VCAM1 NAb. The reduction in Ccr2-RFP+ cells-associated fluorescent pixel area VCAM1 NAb treatment is shown. Unpaired t-test. **, *p*<0.01. ns indicates no significant differences. Each dot indicates a randomly selected ROI (n=11) obtained from mice (n=3 IgG and 4 VCAM1NAb) per group.

To investigate the contribution of endothelial VCAM1 to Mϕ recruitment and MG activation in bAVM lesions, we treated *Cx3cr1-gfp;Ccr2-rfp* mice with a VCAM1 neutralizing Antibody (VCAM1 NAb) starting 5 weeks after AAV-BR1-KRAS^G12V^ injection, with three doses administered at 3-day intervals (**Figure 3E**). We performed intravital imaging before and after treatment to monitor GFP+ Cx3cr1-expressing MG and RFP+ Ccr2-expressing Mϕ within bAVM lesions. In IgG-treated mice, GFP+ cell numbers increased over time, whereas RFP+ cell numbers showed a similar but non-significant upward trend, consistent with progressive myeloid cell accumulation in the bAVM lesions (**Figure 3F**, **3G**, and **3I**). In contrast, VCAM1 NAb treatment markedly reduced RFP+ and GFP+ cell accumulation (**Figure 3F, 3H,** and **3J**). Quantitative intravital imaging further showed that VCAM1 neutralization reduced CCR2+ Mϕ infiltration in bAVM lesion by 64 % (*p*<0.01). These findings provide functional evidence that endothelial VCAM1 contributes to Ccr2+ Mϕ accumulation and modulates local microglial responses in mutant KRAS-induced bAVMs.

### VCAM1 expression is associated with hemorrhage in bAVMs

We previously reported that hemorrhagic conversion in bAVMs is associated with Mϕ/MG accumulation and that depletion of Mϕ/MG or inhibition of Mϕ/MG-associated inflammatory mediators attenuates hemorrhage in bAVMs.^23,32^ To next determine whether endothelial VCAM1 is associated with the Mϕ/MG associated hemorrhage, we compared endothelial VCAM1 expression across bAVM lesions with different degrees of hemorrhage, classified as Ter-119^high^ or Ter-119^low^ based on extravasated red blood cells (RBCs) numbers, at 6 weeks after AAV-BR1-KRAS^G12V^ injection (**Figure 4A** through **4C**). Endothelial VCAM1 expression was significantly increased in bAVM regions containing Ter-119+ RBCs compared with intact regions (**Figure 4B** and **4D**). Notably, endothelial VCAM1 expression was further increased in Ter-119^high^ lesions compared with Ter-119^low^ lesions (**Figure 4D**), which also exhibited greater accumulation of Iba1+ cells (**Figure 4E**). These findings demonstrate an association between increased endothelial VCAM1 expression, greater hemorrhagic burden, and local myeloid cell accumulation in bAVM lesions.

**Figure 4.**
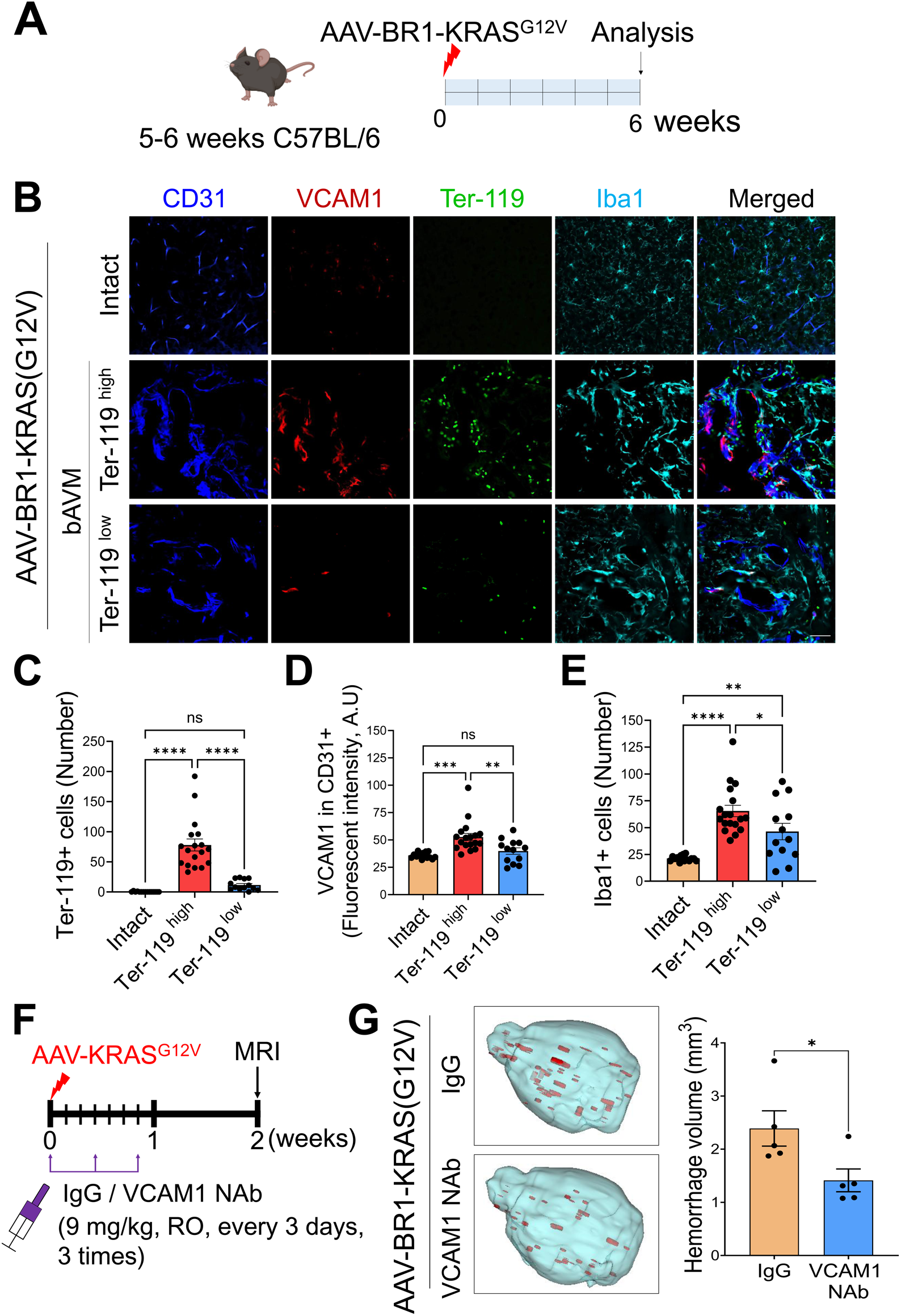
Endothelial VCAM1 expression is associated with hemorrhagic burden in bAVM lesions. **A,** Schematic showing the experimental timeline. C57BL/6 mice received AAV-BR1-KRAS^G12V^ or -AAV-BR1-eGFP (control) and were analyzed 6 weeks after injection. **B,** Representative immunofluorescence images showing VCAM1 expression (red) in CD31+ vessels (blue) in bAVM lesions with high (Ter-119^high^) or low (Ter-119^low^) RBC accumulation. Note that areas with high Ter-119 and VCAM1 expression showed increased accumulation of Iba1+ MG/Mϕ in bAVM lesions. Scale bar = 50 μm. **C** through **E**, Quantification of parenchymal Ter-119+ RBCs (**C**), VCAM1 fluorescence intensity in CD31+ vessels (**D**), and Iba1+ cell numbers (**E**) in intact regions and in Ter-119^high^ or Ter-119^low^ areas of bAVM lesions. ANOVA. *, *p*<0.05, **, *p*<0.01, ***, *p*<0.001, ****, *p*<0.0001. ns indicates no significant differences. Each dot indicates a randomly selected ROI (n=13-18) obtained from mice (n=5-7) per group. **F,** Schematic showing the early VCAM1 neutralization protocol. bAVM mice received VCAM1 NAb starting on the day of AAV-BR1-KRAS^G12V^ injection, followed by two additional doses at 3-day intervals. T_2_-weighted MRI was performed 2 weeks after AAV injection. **G,** Representative MRI-based volumetric images and quantification of hemorrhage volume in control IgG-and VCAM1 NAb-treated bAVM mice. Unpaired t-test. *, *p*<0.05. Each dot indicates the hemorrhage volume in an individual mouse (n=5) per group.

To next investigate whether VCAM1 contributes to hemorrhagic conversion during early bAVM development, we performed a proof-of-concept experiment in which mice received VCAM1 NAb immediately after AAV-BR1-KRAS^G12V^ injection, with three doses administered at 3-day intervals (**Figure 4F**). MRI performed 2 weeks after AAV injection revealed significant reduction in hemorrhage volume in VCAM1 NAb-treated mice compared with IgG-treated mice (**Figure 4G**).

We next investigated whether VCAM1 neutralization could also attenuate hemorrhage at a later stage of bAVM development. VCAM1 NAb treatment was initiated 5 weeks after AAV-BR1-KRAS^G12V^ injection using the same three-dose regimen. In contrast to early treatment, late VCAM1 neutralization did not significantly reduce hemorrhage volume compared with IgG treatment, although a trend toward reduced hemorrhage was observed (**Figure S3A** and **S3B**). Notably, late VCAM1 NAb treatment significantly reduced the numbers of bushy or amoeboid morphology of Cx3cr1-gfp+ MG and Ccr2-RFP+ Mϕ in bAVM lesions compared with IgG treatment (**Figure S3C** through **S3E**). Thus, although VCAM1 neutralization modulated MG/Mϕ responses at later stages, its effect on established hemorrhage was limited under the treatment condition tested. Together with our findings that VCAM1 neutralization reduces Mϕ infiltration and attenuates MG activation, these results suggest VCAM1 contributes to myeloid cell responses during bAVM development and early hemorrhagic progression.

### Genetic reduction of endothelial VCAM1 attenuates Mϕ/MG responses in bAVMs

To further investigate whether endothelial VCAM1 regulates Mϕ recruitment and MG activation in bAVM lesions, we generated mice with endothelial-specific VCAM1 deletion using *Cdh5-CreERT2;VCAM1^fl/fl^* (VCAM1-cKO) mice.^34^ We first assessed the tamoxifen (TM)-induced VCAM1 deletion in VCAM1-cKO mice, we administered TM (1 mg/kg/day) or corn oil for 10 consecutive days beginning at 3 weeks of age. Four weeks after the final injection, we tested isolated ECs from the whole brain by qRT-PCR for Vcam1 and Cdh5 (VE-cadherin). TM treatment reduced VCAM1 mRNA expression without significantly altering Cdh5 mRNA expression (**Figure S4A**).

We next evaluated the effects of TM-induced endothelial VCAM1 deletion using a second defined dosing regimen. TM (40 mg/kg/day) was administered for 5 consecutive days beginning at 4 weeks of age, and AAV-BR1-KRAS^G12V^ was injected 7 days after the final TM injection to induce bAVM formation.^35^ VCAM1-cKO mice exhibited reduced endothelial VCAM1 expression (**Figure S4B** and **S4C**), accompanied by attenuated Mϕ accumulation and MG activation compared with control mice (**Figure S5A** through **S5C**). Further quantification demonstrated reduced numbers of F4/80+ Mϕ or Ly6g+ neutrophils in bAVM lesions following endothelial VCAM1 deletion, supporting a role for endothelial VCAM1 in promoting the accumulation of peripheral myeloid cells within bAVM lesions (**Figure S6A** through **S6E**).

To next examine the effects of sustained endothelial VCAM1 reduction, we used a continuous TM-containing chow regimen. Beginning at 4 weeks of age, VCAM1-cKO mice were fed either regular chow (control) or TM-containing chow to induce bAVM formation. Immunofluorescence analysis confirmed reduced endothelial VCAM1 expression in bAVM lesions of VCAM1-cKO mice compared with controls (**Figure 5A** through **5C**). Importantly, endothelial VCAM1 deletion was accompanied by reduced Mϕ accumulation and attenuated MG activation, as indicated by fewer Iba1+ cells exhibiting bushy or amoeboid morphologies (**Figure 5D** through **5F**). Together, these findings provide genetic evidence that endothelial VCAM1 contributes to peripheral myeloid cell accumulation and MG activation in mutant KRAS-induced bAVMs.

**Figure 5.**
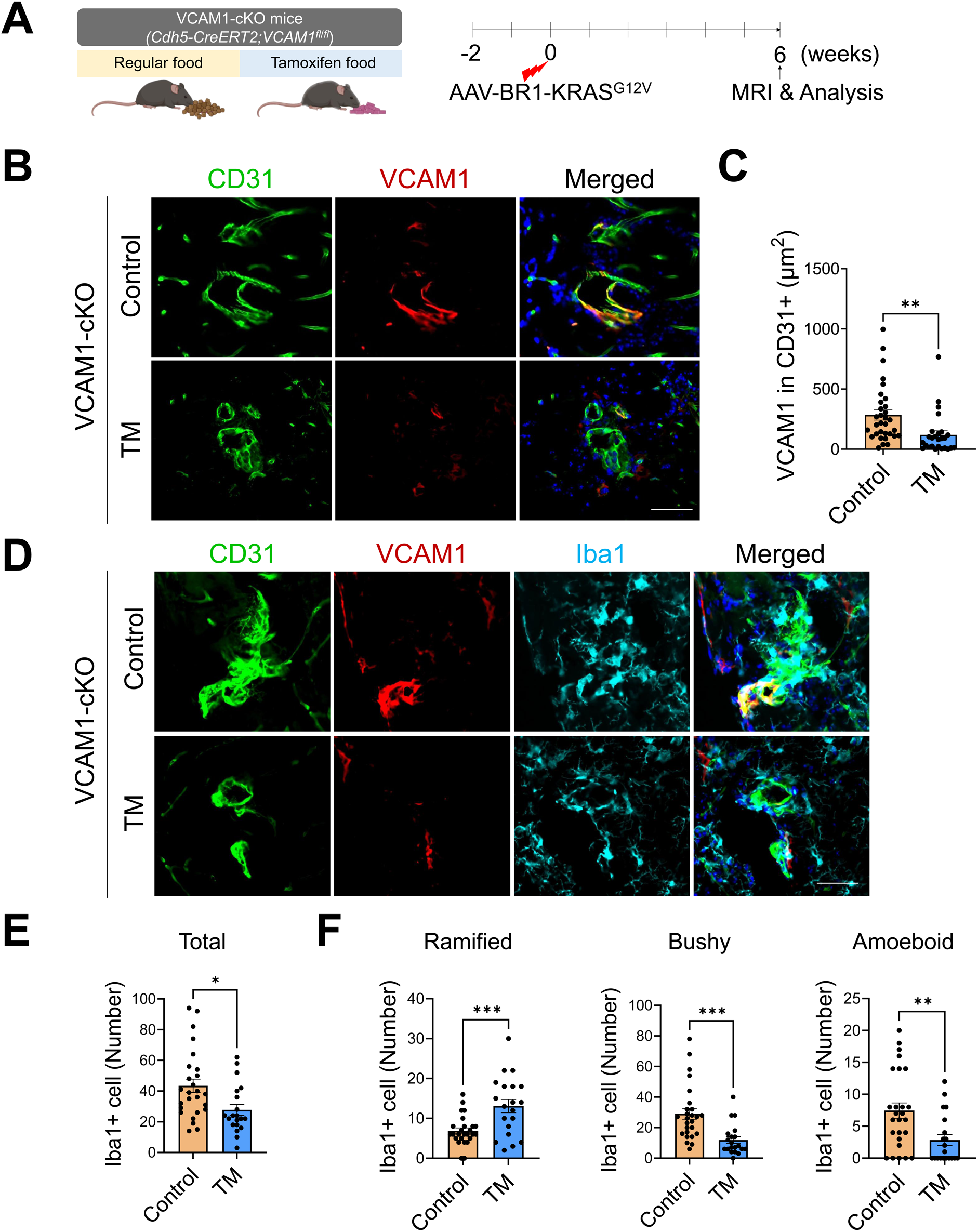
Endothelial VCAM1 deletion reduces MG/Mϕ accumulation and activation-associated morphological changes in bAVM lesions. **A,** Schematic showing the experimental protocol. VCAM1-cKO mice (*Cdh5-creERT2;VCAM1^fl/fl^*) received tamoxifen-containing chow or regular chow (control), starting 2 weeks before AAV-BR1-KRAS^G12V^ injection. Brains were collected 6 weeks after AAV injection. **B,** Representative immunofluorescence images showing reduced VCAM1 expression in CD31+ ECs in bAVM lesions from VCAM1-cKO mice compared with controls. Scale bar = 50 µm. **C,** Quantification of VCAM1+ fluorescence area in CD31-positive vessels from control and VCAM1-cKO mice. Unpaired t-test. **, *p*<0.01. Each dot indicates a randomly selected ROI (n=26-32) obtained from mice (n=8-9) per group. **D,** Representative immunofluorescence images showing Iba1+ MG/Mϕ in bAVM lesions from control and VCAM1-cKO mice. Scale bar = 50 µm. **E** and **F,** Quantification of total Iba1+ cells and the numbers of cells with ramified, bushy, and amoeboid morphologies in bAVM area from control and VCAM1-cKO mice. Unpaired t-test. *, *p*<0.05, **, *p*<0.01, ***, *p*<0.001. Each dot indicates a randomly selected ROI (n=20-26) obtained from mice (n=8-9) per group.

### Genetic reduction of endothelial VCAM1 attenuate hemorrhagic progression in bAVMs

To determine whether endothelial VCAM1 contributes to hemorrhagic progression in bAVMs, we quantified hemorrhage volume in VCAM1-cKO and control mice using MRI. Endothelial VCAM1 deletion significantly reduced hemorrhage volume in mutant KRAS-induced bAVM mice compared with controls (**Figure 6A** through **6C**). We next examined vascular permeability and RBC extravasation to determine whether reduced hemorrhage was associated with improved vascular integrity. Consistent with the reduction in hemorrhage volume, endothelial VCAM1 deletion attenuated vascular leakage, as demonstrated by reduced BSA-647 extravasation and decreased accumulation of Ter-119+ RBCs within and surrounding bAVM vessels (**Figure 6D** through **6H**). Similarly, a separate cohort of mice receiving TM injections showed reduced Ter-119+ RBC extravasation in bAVM lesions (**Figure S7A** through **S7C**).

**Figure 6.**
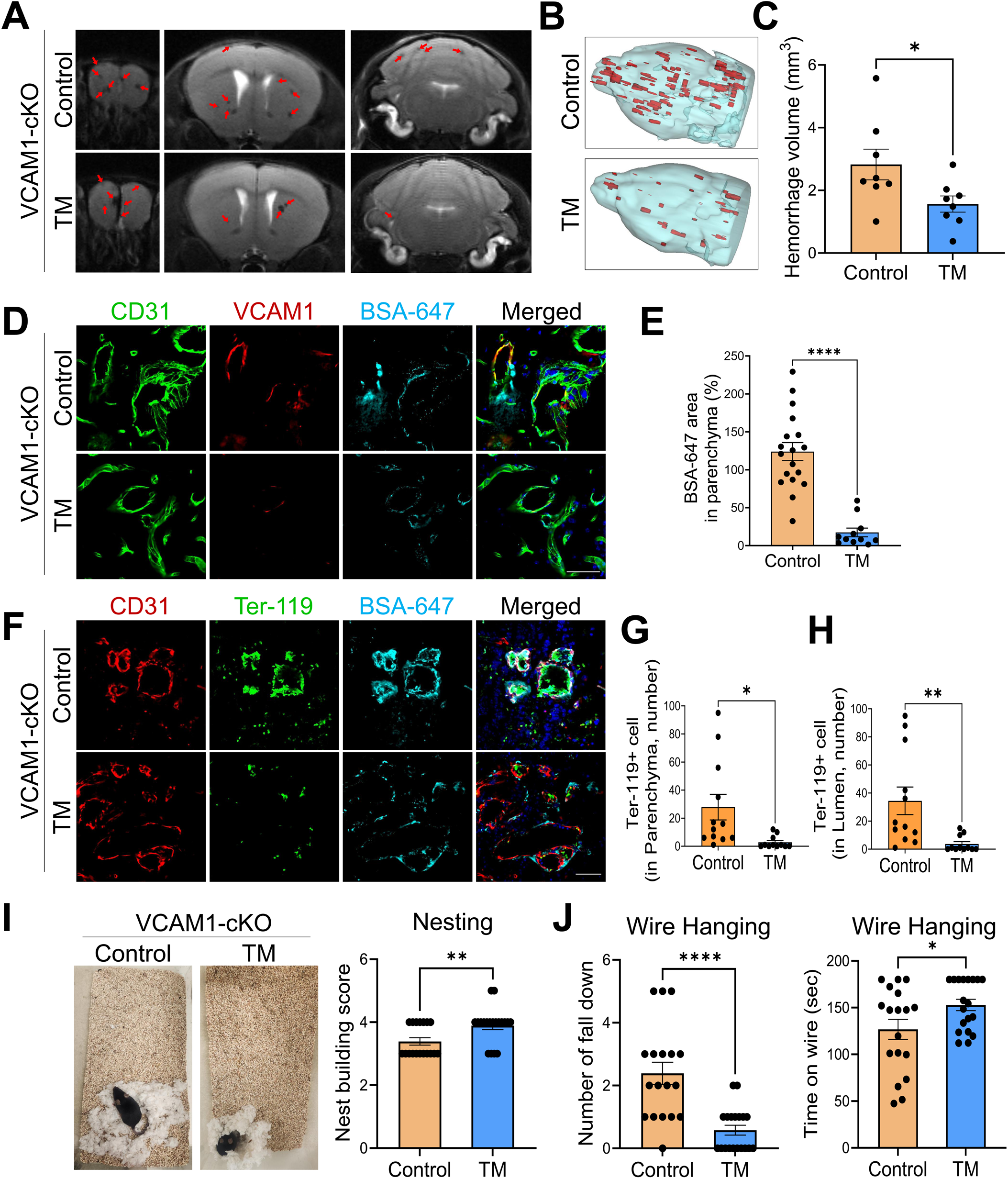
Endothelial VCAM1 deletion attenuates hemorrhagic burden and associated functional deficits in KRAS^G12V^-induced bAVM mice. **A, B,** VCAM1-cKO mice received TM-containing food or regular chow, starting 2 weeks before AAV-BR1-KRAS^G12V^ injection. T_2_-weighted MRI was performed before brain collection at 6 weeks after AAV injection. Representative T_2_-weighted MRI images (**A**) and volumetric images (**B**) showing that endothelial VCAM1 knockdown reduced the ICH lesions in bAVM mice. **C,** Quantification of T2-MRI-defined hemorrhage volume in control and TM-treated VCAM1-cKO mice. Unpaired t-test. *, *p*<0.05. Each dot indicates the hemorrhagic volume of an individual mouse (n=8) per group. **D** through **H,** Representative immunofluorescence images showing endothelial VCAM1 expression (**D**), parenchymal Ter-119+ RBC accumulation (**F**) in bAVM lesions, with BSA-647 extravasation from control-or TM-treated VCAM1-cKO mice. Scale bar = 50 μm. Quantification of BSA-647 extravasation (**E**), parenchymal Ter-119+ RBCs (**G**), and intravascular Ter-119+ RBCs (**H**) in bAVM lesions from control-or TM-treated VCAM1-cKO mice. Unpaired t-test. *, *p*<0.05; **, *p*<0.01. ****, *p*<0.0001. Each dot indicates a randomly selected ROI (n=11-12) obtained from mice (n=5) per group. **I, J,** Behavioral assessment of control-or TM-treated VCAM1-cKO mice. Representative images and quantification of nesting behavior (**I**), which is representative social cognition test and wire-hanging performance, assessed by the number of falls and time spent on the wire (**J**). Unpaired t-test. *, *p*<0.05, **, *p*<0.01, ****, *p*<0.0001. Each dot indicates an individual mouse (n=18-19) per group.

We further evaluated neurological and functional outcomes in these mice. Endothelial VCAM1 deletion was associated with improved behavioral performance, reflected by higher nesting scores and fewer falls during the wire-hanging test (**Figure 6I** through **6J**). Together, these findings demonstrate that endothelial VCAM1 contributes to vascular leakage and hemorrhagic conversion in mutant KRAS-induced bAVMs, and that endothelial VCAM1 deletion reduces hemorrhagic burden and associated functional deficits.

## DISCUSSION

In this study, we identified endothelial VCAM1 as a mediator linking oncogenic endothelial KRAS signaling to myeloid cell accumulation and hemorrhagic progression in bAVMs. Integrated analyses of human transcriptomic datasets, bAVM tissues, and plasma samples demonstrated increased endothelial VCAM1 expression in both unruptured and ruptured bAVMs, accompanied by elevated circulating sVCAM1. In a mutant KRAS-induced bAVM model, endothelial VCAM1 expression was increased and associated with the accumulation of monocyte-derived Mϕ and activation of brain-resident MG. Functionally, VCAM1 neutralization reduced Mϕ accumulation and attenuated MG activation, while endothelial-specific VCAM1 deletion independently reproduced these effects and reduced vascular leakage, hemorrhage progression, and associated behavioral deficits. Together, these findings support a model in which oncogenic KRAS-induced endothelial VCAM1 contributes to myeloid cell responses and vascular inflammation, thereby promoting bAVM instability and hemorrhagic progression.

VCAM1 is a well-established endothelial adhesion molecule that facilitates recruitment and transendothelial migration of monocyte-derived Mϕ, particularly through potential interactions with VLA-4 on monocyte-derived Mϕ.^17^ Our findings extend this established function to KRAS-driven bAVMs, where increased endothelial VCAM1 may facilitate inflammatory myeloid cell accumulation, including brain-resident MG within bAVM lesions. Notably, human bAVM tissue analyses demonstrated increased VCAM1 expression in both unruptured and ruptured bAVMs compared with STA controls, whereas ICAM1 expression was significantly increased only in ruptured bAVM lesions. Consistent with these findings, analysis of a published human endothelial scRNA-seq dataset revealed increased VCAM1 expression in unruptured bAVM ECs compared with control ECs. These observations suggest that endothelial VCAM1 upregulation from unruptured to ruptured is associated with the conversion or progression of bAVM rupture state and is not simply a consequence of overt hemorrhage. Furthermore, within the mouse bAVM model, bAVM lesions with greater Ter-119+ RBC extravasation exhibited higher endothelial VCAM1 expression and greater Iba1+ cell accumulation and hemorrhagic burden. Together these findings link endothelial VCAM1 expression to local myeloid cell accumulation and hemorrhagic burden. Although the mechanism by which oncogenic mutant KRAS induces endothelial VCAM1 remain to be fully determined, previous studies have implicated downstream ERK1/2 signaling in the regulation of VCAM1-dependent leukocyte trans-endothelial interactions.^36^ These pathways warrant further investigation in mutant KRAS-driven bAVMs.

Our pharmacologic intervention studies further suggest that VCAM1 contributes to hemorrhagic progression, particularly during early bAVM development. Consistent with our previous findings that depletion of Mϕ/MG using clodronate liposomes attenuates hemorrhage in mutant KRAS-driven bAVMs,^23^ early VCAM1 neutralization reduced cerebral microbleeds and hemorrhagic volume. In contrast, VCAM1 neutralization initiated at a later stage reduced MG accumulation and attenuated microglial morphological changes, but did not significantly reduce established hemorrhagic burden. These findings suggest that VCAM1-dependent inflammatory responses may be particularly relevant during early bAVM lesion development and hemorrhagic progression, whereas inhibiting VCAM1 later stages may modulate myeloid cell responses without substantially reversing established hemorrhage. This temporal distinction may have clinical relevance because preventing hemorrhage in unruptured bAVMs remains a major unmet need, as underscored by the ARUBA (A Randomized Trial of Unruptured Brain Arteriovenous Malformation) trial. Although our findings provide a rationale for investigating early endothelial inflammatory signaling as a therapeutic target, it remains unclear whether VCAM1-directed intervention can reduce hemorrhagic risk in patients with unruptured bAVMs.

The mechanism by which endothelial VCAM1 influences myeloid cell responses may extend beyond leukocyte adhesion and recruitment. In the present study, both Mϕ and MG expressed VLA-4 components, and ITGA4 expression was increased in bAVM lesions compared with intact regions. These findings establish the presence of the VCAM1-VLA-4 ligand-receptor pair within the bAVM microenvironment and provide a potential mechanism for endothelial-myeloid interactions. However, we did not directly inhibit VLA-4 or genetically manipulate its components; therefore, whether VLA-4 is required for VCAM1-dependent Mϕ recruitment or microglial responses remains to be determined. In addition to its membrane-bound form, VCAM1 can undergo proteolytic shedding to generate sVCAM1, raising the possibility that endothelial VCAM1 may also influence MG through soluble signaling. We found that mutant KRAS increased VCAM1 expression and sVCAM1 release from cultured ECs transfected with mutant KRAS. Furthermore, recombinant sVCAM1 treatment increased the expression of inflammatory mediators, IL-6 and IL-1β, as well as MMP9 and VEGFA in BV2-MG. MMP-9 has been implicated in extracellular matrix remodeling and vascular barrier disruption, whereas VEGFA can increase vascular permeability. These observations suggest that sVCAM1 may contribute to microglial inflammatory and permeability-associated responses.^37^ Notably, these responses resemble those induced by KRAS-EC-CM in our previous study.^32^ However, the extent to which sVCAM1 accounts for the effects of KRAS-EC-CM remains to be established. Whether MG-derived VEGFA or other inflammatory mediators directly contribute to vascular leakage in bAVMs also warrants further investigation.

Several limitations should be further considered. First, the plasma sVCAM1 analysis was limited by the small number of available samples from patients with unruptured bAVMs, reflecting the challenges of obtaining blood samples from this patient population. Nevertheless, our human bAVM tissue analyses independently demonstrated increased VCAM1 expression in unruptured bAVMs. The current plasma data do not establish circulating sVCAM1 as a biomarker for predicting hemorrhagic risk, and larger prospective studies are needed to determine whether circulating sVCAM1 reflects bAVM endothelial activity or correlates with bAVM lesion instability. Second, although our transcriptomic and histological findings demonstrate VLA-4 expression in Mϕ/MG, the functional contribution of VLA-4 to VCAM1-dependent myeloid cell recruitment and activation remains untested. Third, endothelial VCAM1 deletion also reduced Ly6g+ neutrophil accumulation in bAVM lesions, suggesting that VCAM1 may influence the recruitment or accumulation of multiple peripheral myeloid populations. Given the potential contribution of neutrophil-associated inflammation to vascular injury future studies should investigate whether neutrophils participate in VCAM1-dependent hemorrhagic progression. Finally, although our findings implicate sVCAM1 as a potential contributor to microglial inflammatory responses, additional studies are needed to identify the relevant downstream signaling pathways and establish their contributions to vascular injury, including inflammatory mediator production, protease activity, and oxidative stress.^38,39^

In conclusion, our findings identify endothelial VCAM1 as an important component of the inflammatory response associated with mutant KRAS-driven bAVMs. The effects of both pharmacological VCAM1 neutralization and endothelial-specific VCAM1 deletion support a functional role for VCAM1 in myeloid cell accumulation, MG responses, and hemorrhagic progression. Importantly, our findings suggest that endothelial VCAM1 may act as an upstream mediator of the inflammatory vascular response that contributes to hemorrhagic progression, rather than simply serving as a marker of established hemorrhage. Therefore, these findings provide a mechanistic basis for further investigating endothelial VCAM1 as a potential therapeutic target to limit inflammatory vascular injury and hemorrhagic progression in bAVMs.

## Acknowledgments

We gratefully acknowledge our colleagues for their valuable insights and engaging discussions throughout this study and manuscript preparation. We extend our appreciation to the staff at AVM Research Foundation (Ryan-Mission possible) and Ryan & Mark & Sherri Logan family for their unwavering support. The authors thank the Dipaolo and Theaker families for generous donations in support of this work. We thank the technical support from the Cancer Prevention and Research Institute of Texas (CPRIT RP240610).

## Author contributions

J.E.P., H.P., O.M., E.S.P. developed experimental methods, performed experiments, and analyzed the datasets; S.W.K., X.C., Z.Z., E.S.P. Performed and analyzed transcriptome data; J.C., S.Y.J. Performed and analyzed proteomics data; J.E.P., H.P. generated and maintained mice; S.H. established MRI protocol and performed the scan; J.K. provided the plasmid for an EC-specific (AAV-BR1) vector; H.A.C. provided human control plasma; J.E.P., P.R.C., O.M., E.K., E.S.P. prepare human bAVM plasma and human bAVM tissue and analyzed human the data; J.E.P., H.P., E.S.P. quantified imaging data; J.E.P., E.S.P. performed cell culture experiment; J.E.P., J.A., E.S.P. conceptualized study; J.E.P., H.P., E.S.P. designed experiments; J.A., E.K., E.S.P. supported acquiring funding or supported funding for the project; J.E.P., J.A., E.S.P. writing/editing the original draft of the manuscript; J.E.P., H.P., O.M., S.W.K., X.C., Z.Z., J.C., S.Y.J., S.H., J.K., H.A.C., J.A., P.R.C., E.K., E.S.P. edited the final manuscript. And E.S.P. supervised the study.

## Nonstandard Abbreviations and Acronyms

*bAVM*: Brain arteriovenous malformation
*ICH*: Intracerebral hemorrhage
*EC*: Endothelial cell
*VCAM1*: Vascular cell adhesion molecule-1
*ICAM1*: Intercellular adhesion molecule-1
*Mϕ/MG*: Macrophages/Microglia
*KRAS-EC*: EC-carrying mutant KRAS (KRAS^G12V^)
*KRAS^G12V/bEC^* mice: Mice harboring KRAS-EC
*BBB*: Blood-brain barrier
*GFP*: Green fluorescence protein
*RFP*: Red fluorescence protein
*AAV*: Adeno-associated virus
*MRI*: Magnetic resonance imaging
*MRA*: MR angiography

